# Identifying essential genes in *Schaalia odontolytica* using a highly-saturated transposon library

**DOI:** 10.1101/2024.07.17.604004

**Authors:** Joseph K. Bedree, Jacob Bourgeois, Pooja Balani, Lujia Cen, Erik L. Hendrickson, Kristopher A. Kerns, Andrew Camilli, Jeffrey S. McLean, Wenyuan Shi, Xuesong He

## Abstract

The unique epibiotic-parasitic relationship between *Nanosynbacter lyticus* type strain TM7x, a member of the newly identified Candidate Phyla Radiation, now referred to as *Patescibacteria*, and its basibiont, *Schaalia odontolytica* strain XH001 (formerly *Actinomyces odontolyticus)*, require more powerful genetic tools for deeper understanding of the genetic underpinnings that mediate their obligate relationship. Previous studies have mainly characterized the genomic landscape of XH001 during or post TM7x infection through comparative genomic or transcriptomic analyses followed by phenotypic analysis. Comprehensive genetic dissection of the pair is currently cumbersome due to the lack of robust genetic tools in TM7x. However, basic genetic tools are available for XH001 and this study expands the current genetic toolset by developing high-throughput transposon insertion sequencing (Tn-seq). Tn-seq was employed to screen for essential genes in XH001 under laboratory conditions. A highly saturated Tn-seq library was generated with nearly 660,000 unique insertion mutations, averaging one insertion every 2-3 nucleotides. 203 genes, 10.5% of the XH001 genome, were identified as putatively essential.

## Introduction

Members of the *Patescibacteria* phylum (formerly Candidate Phyla Radiation) live as obligate epibionts of host bacteria, making their isolation and characterization difficult. While a number of *Saccharibacteria* and host pairs have been recently isolated (1-4), the *Nanosynbacter lyticus* type strain, TM7x (HMT_952), and *Schaalia odontolytica* (formerly *Actinomyces odontolyticus subsp. actinosynbacter*) strain XH001 co-culture is the first and most well-studied pair (5-17). Studying their interaction has been stymied due to TM7x’s recalcitrance to growth in independent culture and paucity of genetic tools for both species. Transposon and site-specific mutagenesis tools for closely related *Saccharibacteria* spp. were recently developed, however (4). Additionally, we developed a site-specific mutagenesis methodology for the basibiont, XH001 (8), which elucidated AI-2 quorum sensing genetic mediators upon TM7x infection, that was built upon the work of *Yeung* et al., who pioneered genetic system development in closely related *Actinomyces* spp. A broad-host-range vector (18), pJRD215 (19), as well as integration vectors were developed for *Actinomyces naeslundi* (20-22). Marker-less, in-frame deletion mutants were developed for *Actinomyces oris* MG-1 using GalK counterselection (23). Additionally, lytic and temperate bacteriophage were used for transfection in *Actinomyces*, however, none were found to infect *Schaalia odontolytica* (22). Most recently, newer genetic systems have been built for *Actinomyces oris* MG-1 through leveraging marker-less, in-frame mutations using fluorescence (mCherry) counterselection (24) or transposon mutagenesis (25). This cumulative knowledge laid the groundwork for developing a high-throughput mutagenesis tool for the study of XH001 (8, 26), which has been a barrier identifying essential genes in *Schaalia* spp. even though such studies have been conducted for the major oral pathogens *Aggregibacter Actinomycetecomitans* (27), *Fusobacterium nucleatum (28, 29), Porphorymonas gingivalis (30)*, and *Streptococcus mutans* (31, 32). Tn-seq (33) is a robust technique for elucidating the essential genes of any bacterium. High-throughput transposon mutagenesis achieves an insertional saturation allowing a probabilistic assessment of essentiality based upon relative gene size and number of insertions (30, 34). The construction of an XH001 Tn-seq library enabled an analysis of essential genes in XH001 and marked a crucial advancement in XH001 genetic tools for the broader scientific community.

## Materials and Methods

### Bacterial Strains, Plasmids, and Media

All bacterial strains, plasmids and media used in this work are listed in Table S1. All growth conditions used followed previous culturing protocols (8), except XH001::Tn*5* mutants were selected on Brain-Heart Infusion (BHI) broth or agar (Difco Laboratories, Detroit, Michigan) with 500 µg/mL of kanamycin sulfate (Fischer Bioreagents, Hampton, NH, United States). All strains whether grown in broth or on agar were incubated at 37°C in microaerophilic conditions (2% O2, 5% CO_2_, balanced with Nitrogen) without shaking using a Whitely Workstation A35 (Microbiology International, Frederick, Maryland).

### Transposon Mutagenesis and Library Collection

XH001 underwent electrocompetent cell preparation and transposition with the EZ-Tn*5* transposon as previously described (16, 25, 35, 36) with the following modifications: electrocompetent cells underwent a glycine shock (5% v/v final concentration) prior to harvesting, which has been previously shown to increase transformation efficiency (25). The transposome mixture was generated by amplifying the EZ-Tn*5* transposon with primers ME-9F and ME-9R from the pMOD-2/Kan215 plasmid as previously described (25), which contained ∼200 ng of transposon in each reaction. 200 *µ*L of electrocompetent cells were transformed per reaction and recovered in 1 mL of BHI at 37 °C for 3 hours. The entire 1.2 mL culture of recovered transformants were plated, 50 *µ*L at a time, to optimize cell spread and were selected on BHI with 500 *µ*g/mL of kanamycin sulfate for up to 7 days in microaerophilic conditions described above. The resulting colonies from each library were pooled with an inoculation loop as previously described (37) and collected in 10 mL of fresh media supplemented with 500 *µ*g/mL of kanamycin sulfate. Six separate libraries were generated, each comprised of three independent transformations to maximize the number of unique EZ-Tn*5* insertions into the chromosome. These individual libraries were aliquoted (∼2.0 × 10^8 CFUs/mL) and stored in 30 % glycerol at -80 °C. One cryovial from each library was later used for DNA isolation.

### DNA Isolation and Deep Sequencing

DNA isolation was performed using Epicentre MasterPure™ Complete DNA and RNA Purification Kit (Lucigen) following the manufacturer’s protocol with the following modifications: pre-lysis incubation of samples with Lysozyme (50 mg/mL) and Mutanolysin (2 mg/mL) at 37°C and included an RNAase step for 30 minutes at 37°C. Library preparation was performed as previously described (30, 38) with the following modifications: during the first PCR amplification step, forward primer 1 was added to a reverse primer mixture consisting of primer 2 and 3. This reverse primer reaction mixture consisted of primer 2, a truncated version of primer 3, which was added at a 10:1 volumetric ratio (primer 2:primer 3, respectively) to enhance amplification and avoid primer dimer formation initially observed in PCR reactions with only primer 1 and primer 3. The second PCR amplification step, the barcode addition, was completed using primer 4 and barcode primers (BC1-12) listed in Table S1. All PCR reactions were completed using Q5® Hot Start High-Fidelity DNA Polymerase (New England Biolabs, Ipswich, MA) following the manufacture’s protocol. For deep sequencing on Illumina’s HiSeq 2500 platform, a custom sequencing primer was used.

### Bioinformatic Analysis for Gene Essentiality

Bioinformatic analysis of the Tn-seq library sequencing datasets was performed as described (34, 39), which first comprised of trimming off the 3’ poly-C adapter and 5’ Ez-Tn*5* Mosaic End (ME) sequences from any given read (sequence) using stringent parameters for cutadapt.py (40). All trimmed reads from each of the six libraries were merged into one file to determine aggregate insertional density using parameters from the previously established (34) approach. The aggregate hopcount analysis was performed with the following parameters: (1) minimum read positional cutoff of 15 (15 reads at a given position needed for analysis); (2) when aggregating EZ-Tn*5* hops into the genome, did not count reads in the first or last 5% of the CDS; (3) only count reads that align perfectly to the reference across the entire read (no mismatches or gaps). For essential gene analysis, the following criteria were used: Dval genome value (defined as the number of reads of each gene divided by the expected number of reads based on gene size) <= 0.01 (34) and minimum region length = 100 base pairs, generating a putatively essential gene list (Table S2). All designated CDS gene loci were then assigned to Clusters of Orthologous Groups as previously described (41, 42).

Spearman correlation matrix comparing DvalGenome values between the 6 individual EZ-Tn*5* mutagenesis libraries assessing all regions within the XH001 genome (intergenic and CDS) was performed using Graph Pad Prism 8 software. All 203 putatively essential genes that we identified (Fig. 2) were visualized using Circos 0.69-9 (https://circos.ca/) (43). All genes were designated by gene name or APY gene loci tags in the light blue outermost ring. Genome insertion snapshots used for highlighting putatively essential genes (Fig. 3, 4) were taken by displaying transposon insertions along the *Schaalia odontolytica* strain XH001 (LLVT01000001.1) genome in Integrated Genome Viewer (https://igv.org/) (44). Putatively essential protein functional domains were identified for XH001 proteins by searching for non-uniform distribution of transposon insertions in candidate genes. After identifying candidate genes, putative homologous protein functional domains for encoded proteins were identified using InterProScan (https://www.ebi.ac.uk/interpro/result/InterProScan/) (45) and plotted with transposon insertions in Integrated Genome Viewer (Fig. 4). COG analysis (Fig. 5) was performed as previously described (42) and figures were generated using Graph Pad Prism 8 software.

## Results

### Mutant Library Construction and Essential Gene Analysis by Tn-seq in XH001

To construct a highly saturated Tn-seq library, the EZ-Tn*5* mini-transposon was selected for its ease of use and its stable, random insertions (47, 48) relative to the Himar1-mariner transposon, which has a strong AT-nucleotide insertional bias (33, 49) that is unsuitable for high GC% organisms. While the original Tn*5* normally shows low efficiency (50), a combination of a hyperactive triple mutation of the Tn*5* transposase (47) and inverted repeat ends containing a mosaic end (ME) sequence (51), increased the transposition insertion efficacy by several orders of magnitude (50). Furthermore, it has been used in several bacterial species (25, 34, 52, 53). The kanamycin-resistance cassette from pJRD215 (19) was cloned into the MCS of the commercial EZ-Tn*5* pMOD vector, yielding a custom EZ-Tn*5* transposon previously demonstrated high rates of transposition in a closely related *Actinomyces* spp. (25). This amenable transposon system was utilized to generate a highly saturated insertion library in XH001 (Fig. 1). A total of six separate mutant libraries were generated (25) and collected (37) using established protocols with modifications (see Methods). Each mutant library underwent DNA extraction, library preparation, and deep sequencing as previously described (30, 38).

**Fig. 1.**
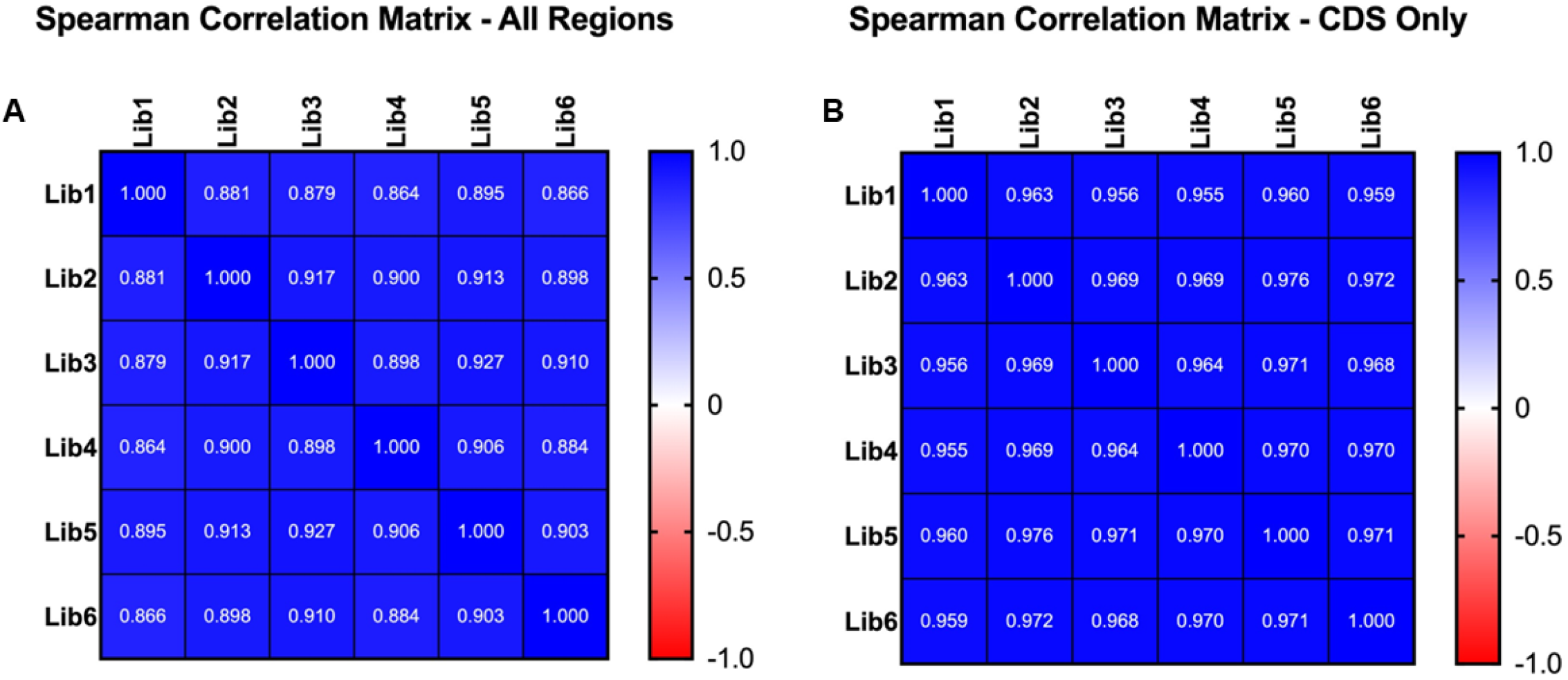
Robust EZ-Tn*5* mutagenesis of the XH001 genome. Fig. 1A represents the Spearman correlation matrix comparing DvalGenome values between 6 individual EZ-Tn*5* mutagenesis experiments assessing all regions within the XH001 genome (intergenic and CDS). Fig. 1B represents the Spearman correlation matrix comparing the 6 individual EZ-Tn5 mutagenesis experiments assessing all CDS regions within the XH001 genome. All correlations were significant (>0.8).

**Fig. 2.**
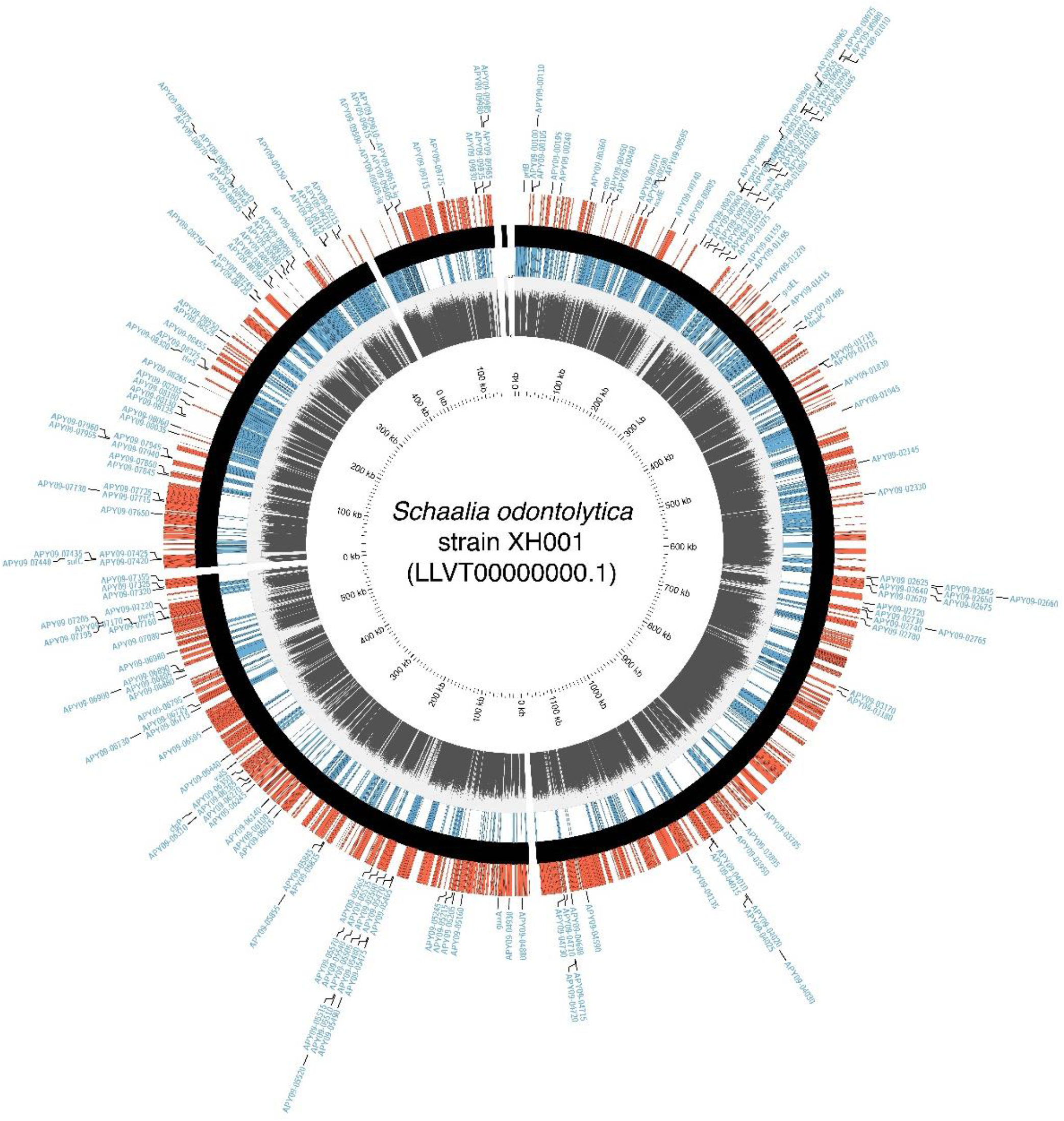
Mapping The Putatively Essential Genes in XH001 for Growth on Solid Media. The total insertion density is log scaled from 10^0^-10^5^ in the grey innermost ring and by strand in the red (+) and dark blue (-) rings. All 203 putatively essential genes are designated by gene name or APY gene loci tags in the light blue outermost ring. This Fig. was generated using Circos 0.69-9 (https://circos.ca/) (43).

**Fig. 3.**
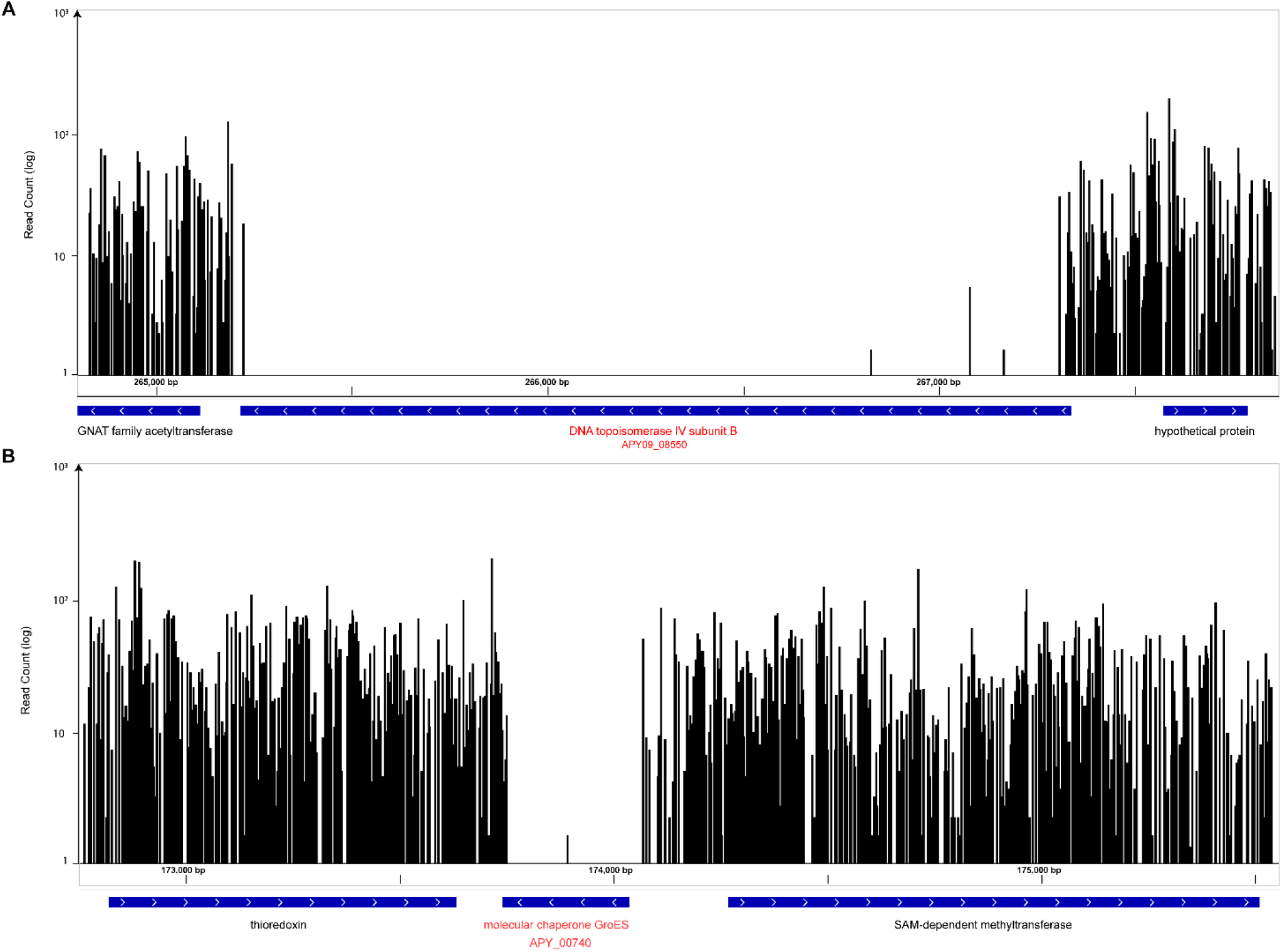
Examples of transposon insertions into highly saturated genes and identification of putatively essential genes. Genome insertion snapshots were taken by displaying unique transposon insertions as a WIG file along the *Schaalia odontolytica* strain XH001 (LLVT01000001.1) genome in Integrated Genome Browser (https://bioviz.org/) (46). Here, genes are displayed as blue arrow boxes in the direction of coding sequence orientation where black bars represent genomic position and number of sequenced insertions. The genes that encode and DNA topoisomerase IV subunit B (panel A) and molecular chaperone GroES (panel B) have almost zero insertions along their coding sequences suggesting they are likely to be essential in this growth condition. Moreover, there is a high density of insertions immediately upstream and downstream of these essential genes, signifying non-essential regions.

**Fig. 4.**
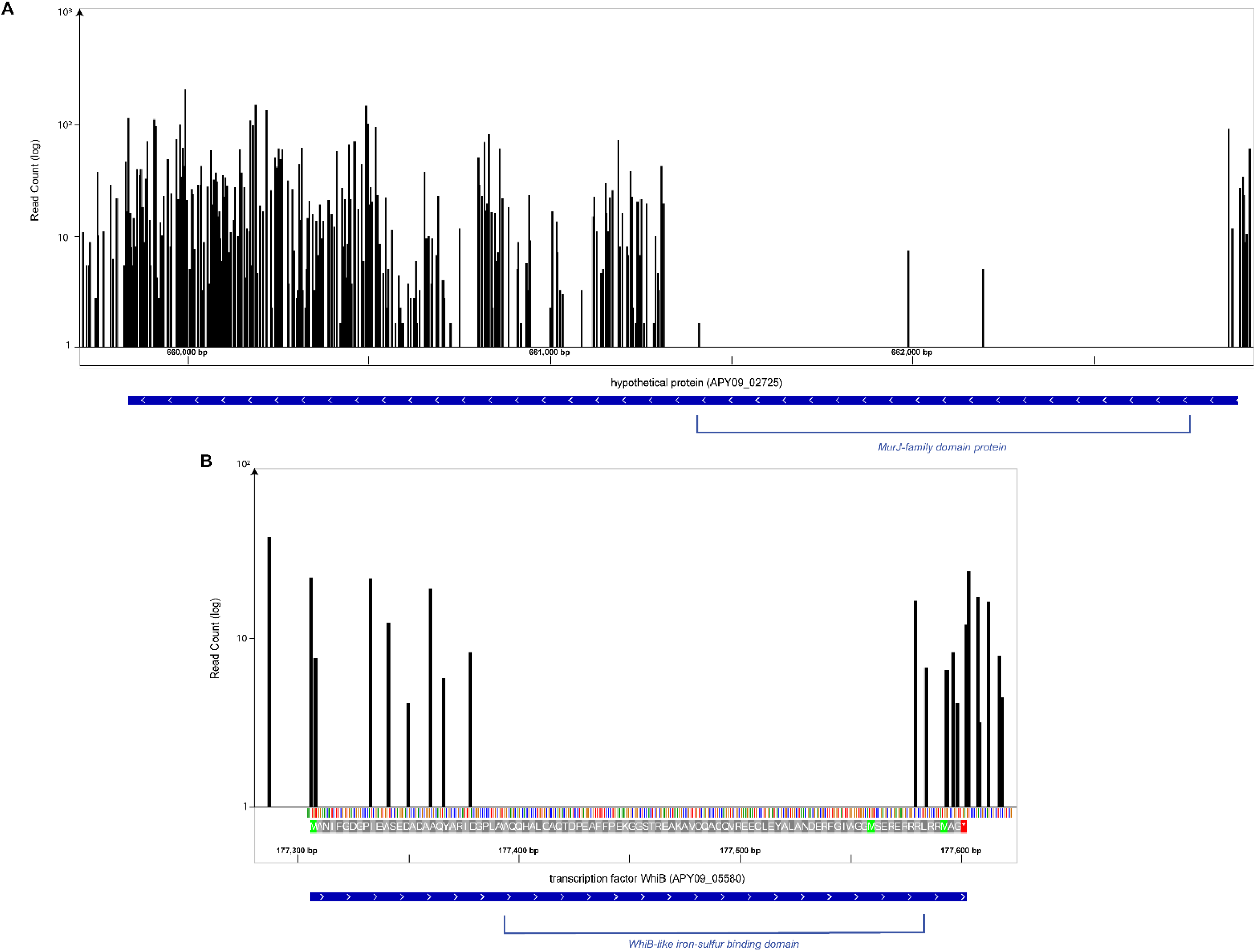
Examples of transposon insertions revealing putative essential protein domains. Non-uniform distribution of transposon insertions in genes suggests the presence of essential protein domains. Putative homologous protein functional domains were identified for XH001 proteins using InterProScan (*https://www.ebi.ac.uk/interpro/result/InterProScan/*) (45). Locus APY09_02725 encodes a putative protein with several unique insertions excluding a region predicted to contain a MurJ-family protein domain (panel A). Transcription factor WhiB (APY09_05580) lacks insertions in its predicted sulfur-binding domain (panel B).

**Fig. 5.**
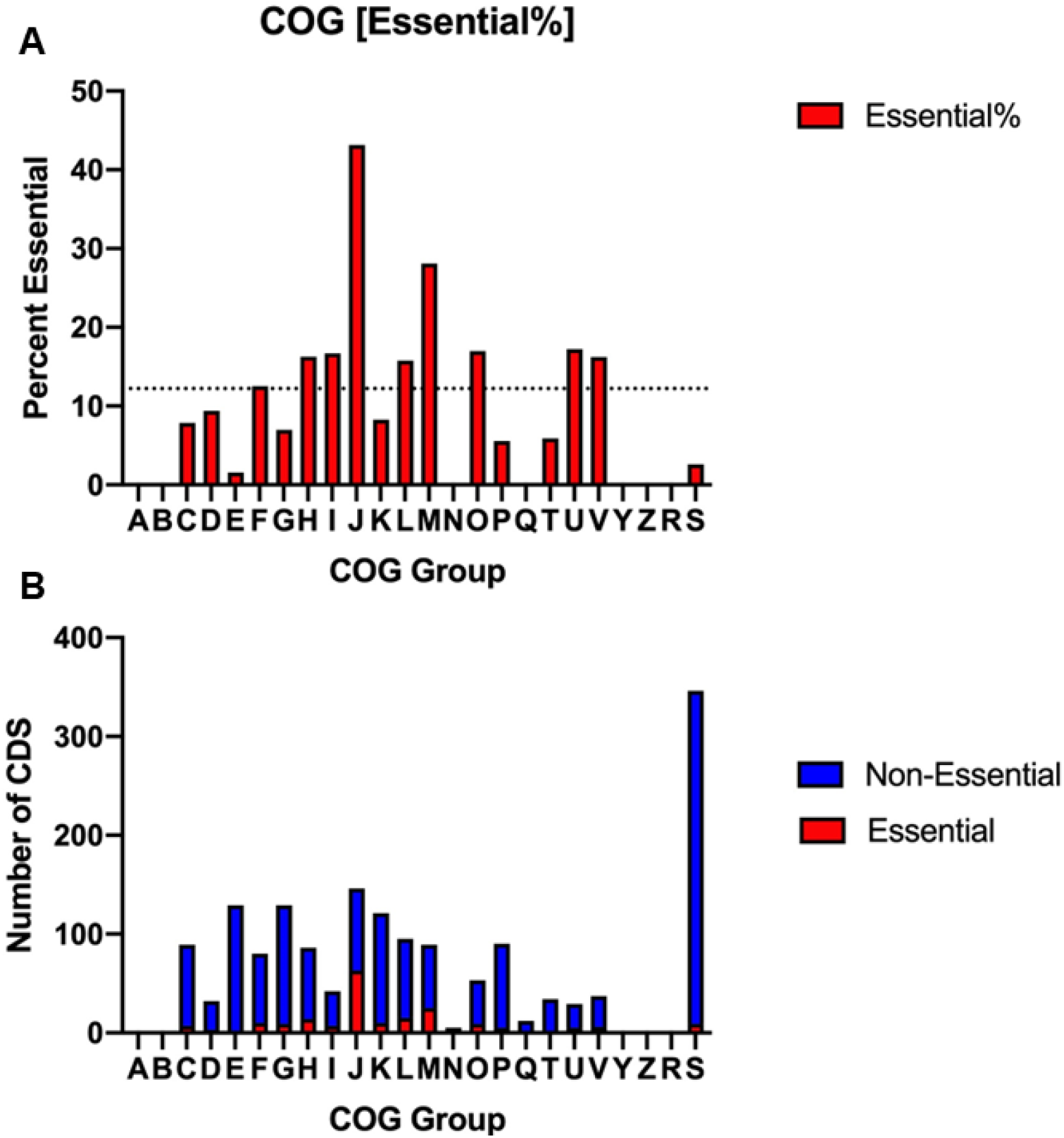
Clusters of Orthologous Groups in XH001. Fig. 5A shows the relative distribution of putatively essential genes in Clusters of Orthologous Groups (COGs). Each bar represents the relative percent of putatively essential genes that lie within that COG, the dotted line at 12.2% represents the ratio of putatively essential genes with known COG designation to all coding genes in XH001 with COG designation. Fig. 5B represents the ratio of putatively essential to non-essential genes with COG designation within each COG group. Both Figs. 5A and B utilize the following X-axis key as described: [A = RNA processing and modification; B = Chromatin structure and dynamics; C = Energy production and conversion; D = Cell cycle control, cell division, and chromosome partitioning, E = Amino acid metabolism and transport, F = Nucleotide transport and metabolism, G = Carbohydrate transport and metabolism, H = Coenzyme metabolism and metabolism, I = Lipid metabolism and metabolism, J = Translation ribosomal structure and biogenesis, K = Transcription, L = Replication, recombination, and repair, M = Cell wall/membrane/envelop biogenesis, N = Cell motility, O = Post-translational modification, protein turnover, chaperone functions; P = Inorganic ion transport and metabolism; Q = Secondary metabolites biosynthesis, transport, and catabolism; T = Signal transduction mechanisms; U = Intracellular trafficking, secretion, and vesicular transport; V = Defense Mechanisms; W = Extracellular structures; Y = Nuclear structure; Z = Cytoskeleton; R = General functional prediction only; S = Function unknown].

### Analysis of location and patterns of insertion sites within a highly-saturated transposon library reveals insights into the genome

The presence of insertions in a gene, which reflects mutant bacterium in which transposons were inserted in those positions, indicates the gene is dispensable in the library BHI growth condition. Likewise, the absence of insertions in a gene suggests that the gene is essential in this growth condition. However, in the absence of additional experiments to prove essentiality on a gene-by-gene basis, we refer to these as putatively essential genes in this study. To identify putatively essential genes in the XH001 genome, we aggregated the transposon sequencing data across six biological replicates to eliminate technical variation by following previously described essential gene parameters cutoffs (30). To evaluate our hypothesized, unbiased insertional mutagenesis with the EZ-Tn*5* transposon, we used Spearman’s rank correlation coefficient to compare the DvalGenome values between all 6 libraries, which revealed consistent insertion saturation results across the replicates (Fig. 1, Table S1). Sequencing showed extensive genome coverage with nearly 660,000 unique insertions, averaging one insertion every 2-3 nucleotides. We then applied the following subsequent analysis criteria: First, poor quality reads (MAPQ < 1) are not used in the analysis. Next, there must be at least 15 sequenced reads for a position to be counted as an insertion. This was done to reduce false insertion sites, which can occur due to sequencing errors, especially with nearly identical repeated sequences in the genome. Finally, as insertions in the extreme start or end of a predicted coding sequence are more likely to yield a functional protein, only insertions after the first 5% or before the last 5% of a gene are counted as insertions within a gene. This stringent selection revealed 203 putatively essential genes (out of 1,936 gene coding sequences) that had zero insertions, of which 10 have no clear orthologue by BLASTp within the Database of Essential Genes (54, 55). This Tn-seq dataset also displayed no insertions in two intergenic regions (Fig. 2, Table S2). Altogether, 10.5% of the XH001 genome is predicted to be essential in the BHI growth condition, consistent with previous reports (30, 52, 56-61).

In a highly saturated transposon library, genes with several unique insertions distributed along the coding sequence are deemed non-essential for the growth condition, whereas genes with no insertions are likely to be essential. Genes that are nearly essential, i.e., that result in a severe growth defect, may have few or no insertions. Moreover, transposon insertions in a gene may exert a polar effect on a downstream essential gene or genes, making it difficult to conclude that the first gene is essential. Therefore, genes that lack insertions within a Tn-seq screen are more correctly referred to as putatively essential. Two examples of genes identified as putatively essential in this study were the genes for chaperone protein DNA topoisomerase IV subunit B (APY09_02670) and GroES (APY09_00740) (Fig. 3). GroES is a chaperonin that is essential in *Escherichia coli* (62). Examination of insertions around *groES* revealed a complete absence of insertions within the coding sequence (Fig. 3). The degree of sequencing depth and insertion complexity argues against random chance as an explanation for lacking insertions within *groES*. Furthermore, using an aggregated library reduces the probability of observing a lack of insertions due to technical error. Finally, the presence of several unique insertions in the flanking genes rules out the possibility that the region in question is a “cold” spot for transposon insertion, further arguing that the lack of insertions in the coding sequence are due to essentiality. DNA topoisomerase IV subunit B (APY09_02670), which encodes a protein essential for chromosomal segregation, showed a similar pattern (Fig. 3).

In addition to identifying putatively essential genes, a highly-saturated transposon insertional library reveals information about additional genetic elements such as promoter regions, protein functional domains, mis-annotations, and operon structure. Specific domains within a protein may be essential even if the overall protein is not, which may be detected by searching for non-uniform distributions of insertions within coding sequences (Fig. 4). For example, examination of insertions within the coding sequence of *whiB* (APY09_05580) revealed a lack of insertions in part of the coding sequence. The WhiB protein is a transcription factor that regulates several functions in actinobacteria (63, 64). Examination of the essential protein domain revealed a predicted sulfur-binding domain with the C-X-X-C motif characteristic to WhiB-family proteins (64, 65). Similarly, analysis of coding sequence APY09_02725 revealed a non-uniform distribution in which the first half of the coding sequence lacks any insertion. Notably, protein domain prediction software suggests this area is homologous to MurJ-family proteins, which are involved in peptidoglycan synthesis and regulation (66). This may suggest that this locus encodes a protein involved in cell wall regulation, encodes a multi-functional protein, represents a fusion protein between an essential MurJ-family protein and a non-essential protein, or may be a mis-annotation of the genomes. Altogether, examination of insertion distribution both within and between predicted coding sequences yields valuable data about genetic structure and protein function.

The *S. odontolytica* strain XH001 genome used in our analysis has a 1.16 Mb genome and contains 1,998 genes of which 1,936 are predicted to be protein-coding (11) (GenBank accession: LLVT01000001.1). The functional distribution of the putatively essential genes was determined by assigning genes to Clusters of Orthologous Groups (COGs) as described (42). As expected, a greater percentage of non-essential genes could not be assigned to COGs, resulting in a higher percentage of putatively essential genes when only examining COGs (12.2%) (Fig. 5A, Table S2). Significant enrichment was seen in ‘H’ (coenzyme metabolism and metabolism), ‘I’ (lipid metabolism and metabolism), ‘J’ (Translation ribosomal structure and biogenesis), ‘L’ (replication, recombination, and repair), ‘M’ (cell wall/membrane/envelop biogenesis), ‘O’ (post-translational modification, protein turnover, chaperone functions), and ‘U’ (intracellular trafficking, secretion, and vesicular transport). Enrichment in this case was simply defined as a proportion of essential genes in a COG group higher than the ratio of essential COG to nonessential COG. This suggests that the core, essential part of the genome focuses on these tasks (i.e., lipid metabolism) rather than the other tasks (i.e., amino acid metabolism). Unexpectedly, ‘V’ (defense mechanisms), was also enriched, potentially suggestive of essential stress responses to the microaerophilic conditions during XH001 culturing. Interestingly, approximately ∼20% (346) of the total CDS within XH001’s genome are assigned to category ‘S’ (function unknown).

## Discussion

Performing essential gene identification studies are extremely important for understanding both bacterial physiology and metabolism in natural biological contexts. Here, we present an essential gene study in a *Schaalia* spp. grown on BHI agar (solid media) in microaerophilic conditions. From this, we identified 203 putatively essential genes representing 10.5% of the XH001 genes. Unsurprisingly, the putatively essential genes had significant enrichment in basic biological functions such as lipid metabolism, replication, and ribosomal structure. However, 10 of these genes, have no clear orthologue by BLASTp within the Database of Essential Genes (54, 55) or no known function (Fig. 5). Moreover, of the non-essential genes in XH001, 20% lacked known assigned function. This highlights major challenges in understanding genetic requirements and biology of microorganisms and is not unique to *Schaalia* spp., since even in the model organism *E. coli*, ∼35% of its genome remains to be understood in terms of gene function (67). In this study, as with any study using insertional mutagenesis, the possibility exists of having no insertions in a particular gene simply because they are polar on a downstream essential gene, yielding false positive identification of the gene as putatively essential. Future work would benefit from individual genetic dissection of these putatively essential or non-essential genes. Together, these data and the XH001 Tn-seq platform presented in this study add a powerful new tool to study *S. odontolytica* XH001 biology and facilitate a better mechanistic understanding of the episymbiotic relationship between XH001 and its epibiont TM7x.

## Acknowledgments

The authors would like to thank Dr. Chenggang Wu for generously providing technical assistance in implementing a viable transposon mutagenesis strategy and the pMOD-2/Kan215 vector. The authors would also like to thank Dr. Javier Fernandez Juarez for Tn-seq library preparation optimization and the Tufts Genomics Core manager, Dr. Albert Tai, for assistance in Tn-seq sequencing and troubleshooting. Research reported in this study was supported by the National Institute of Dental and Craniofacial Research of the National Institutes of Health under Award Numbers F31DE026057, KL2TR002317, R01DE023810, R01DE030943 and R01DE029479.

## Notes

### Competing Interest Statement

The authors have declared no competing interest.

